# Experience-dependent development and maintenance of binocular neurons in the mouse visual cortex

**DOI:** 10.1101/614768

**Authors:** Kyle R. Jenks, Jason D. Shepherd

## Abstract

The normal development of neuronal circuits requires both hard-wired gene expression and experience. Sensory processing, such as vision, is especially sensitive to perturbations in experience. However, the exact contribution of experience to neuronal visual response properties and binocular vision remains unknown. To determine how visual response properties develop *in vivo*, we used single cell resolution two-photon calcium imaging of mouse binocular visual cortex at multiple time-points after eye opening. Few neurons are binocularly responsive immediately after eye opening and respond solely to either the contralateral or ipsilateral eye. Binocular neurons emerge during development, which requires visual experience, and show specific tuning of visual response properties. As binocular neurons emerge, activity between the two eyes becomes more correlated in the neuropil. Since experience-dependent plasticity requires the expression of activity-dependent genes, we determined whether the plasticity gene *Arc* mediates the development of normal visual response properties. Surprisingly, rather than mirroring the effects of visual deprivation, mice that lack *Arc* show increased numbers of binocular neurons during development. Strikingly, removing *Arc* in adult binocular visual cortex increases the numbers of binocular neurons and recapitulates the developmental phenotype, suggesting cortical circuits that mediate visual processing require ongoing experience-dependent plasticity. Thus, experience is critical for the normal development and maintenance of circuits required to process binocular vision.

## INTRODUCTION

Sensory processing requires the development of neuronal circuits that can adapt to environmental stimuli. Thus, the shaping of brain circuits requires both hard-wired patterning and experience-dependent plasticity. These experience-dependent processes occur early in development during critical periods of heightened plasticity. This is particularly evident in the visual cortex (V1), where manipulation of visual experience early in life can lead to dramatic changes in function and structure^1^. Closing one eye for several days drives the loss of input from the deprived eye in V1, resulting in ocular dominance (OD) plasticity. While the mechanisms of OD plasticity are well studied^2^, it remains unclear whether similar mechanisms mediate the development of normal visual response properties.

In most mammals, V1 receives and integrates visual input from both eyes. Binocular V1 needs to integrate inputs from the two eyes to properly resolve an accurate representation of visual space. In cats and most primates, which share forward facing eyes and large binocular fields, inputs between the two eyes are arranged in alternating OD columns. These columns appear perpendicular to orientation columns that correspond to the orientation of edges (dark-light borders) in the neuron’s receptive field and are dominated by the contralateral eye^3^. This structural organization seems to arise independent of visual experience, although other visual response properties, such as direction selectivity seem to require experience^4–6^. Mice have a smaller binocular visual field than cats and primates, and lack OD columns. However, neurons within mouse binocular V1 still display a range of orientation selectivity and a bias towards the contralateral eye^7^. Recent studies have also uncovered spatial organization of orientation selectivity as a function of cortical distance and cortical depth, consistent with primitive columnar organization^8^.Studies using intrinsic imaging in mouse V1 suggest that eye-specific plasticity begins at eye opening, especially in the establishment of ipsilateral connectivity^9^. Initial input into V1 is mostly driven by the contralateral eye and the refinement of ipsilateral eye responsiveness requires patterned contralateral visual input, although it is unclear how this large-scale refinement of ipsilateral input relates to integration of binocular input at the level of single neurons.

OD plasticity is mediated by molecular mechanisms reminiscent of cellular models of plasticity such as long-term potentiation and depression (LTP/LTD)^10^. However, little is known about the contribution of these mechanisms to the development and refinement of normal visual response properties. The immediate early gene *Arc* is induced by visual experience^11^ and is necessary for OD plasticity and LTD in binocular V1. Arc expression declines past the close of the critical period and increasing Arc expression in the adult brain can restore juvenile like OD plasticity^12,13^. Arc has also been implicated in the tuning of orientation selectivity^14^. Thus, Arc may also play a critical role in experience-dependent development of visual response properties.

To determine how visual response properties develop, we used two-photon *in vivo* calcium imaging in mouse binocular V1 to examine visual response properties of individual neurons at various points early after eye opening. In order to determine the contribution of experience and synaptic plasticity, we also imaged dark reared mice and mice where Arc expression was manipulated.

## RESULTS

### Visually responsive neurons emerge after eye-opening

To measure visual response properties of single neurons, we conducted acute, two-photon calcium imaging through cranial windows implanted over binocular V1 (Figure 1A) in cohorts of wild type mice expressing GCaMP6s driven by the thy1 promoter in excitatory neurons of layer 2/3 (L2/3)^15,16^. To capture changes in response properties after eye-opening, we imaged at three points in early development (Figure 1B): postnatal day (P)14 corresponding to just after eye opening, P20 immediately prior to the start of the canonical OD plasticity critical period, and P30, the peak of the critical period. We presented visual stimuli (drifting, sinusoidal grating from 0-330° in 30° increments), interleaved with grey screen presentations, to each eye independently in pseudorandom order while recording in the same hemisphere. Regions of interest (ROIs) corresponding to neuronal soma were selected using the program Suite2P^17^ and manually annotated (see methods). We subtracted the neuropil fluorescence (see methods) around the soma to remove contaminating signal from surrounding processes (Figure 1C, left panel)^18^. Neuronal activity was Z-scored using the mean and standard deviation of activity during the grey screen presentations (Figure 1C, Right panel). Cells were considered visually responsive if the mean response amplitude (from 0-5 second post stim onset) to a visual stimulus was 0.5 standard deviations above the mean of the responses to the grey screen, and if the responses to individual presentations of that stimulus were significantly larger in mean amplitude (p<=0.05, paired *t*- test) than the preceding grey screen periods. Visually responsive neurons were further classified as either binocular, where the neuron showed significant visual responses to stimulation of both the contralateral and ipsilateral eye independently, or monocularly responsive if they responded solely to either the contralateral eye or ipsilateral eye (Figure 1D). The spatial distribution of binocular and ipsilateral responses appeared widely dispersed, discounting the possibility that our recordings were in or near the boundary of monocular V1. The percent of visually responsive neurons increased significantly, from 13±3% (mean±standard error) at P14 to 39±9% at P30 (Figure 1E, wild type (WT) P14=27±7 total responsive neurons/mouse N=5; P20=59±7 neurons/mouse N=5, P30=60±3 neurons/mouse N=6; F_2,13_=30.8, p<0.0001, ANOVA).

**Figure 1:**
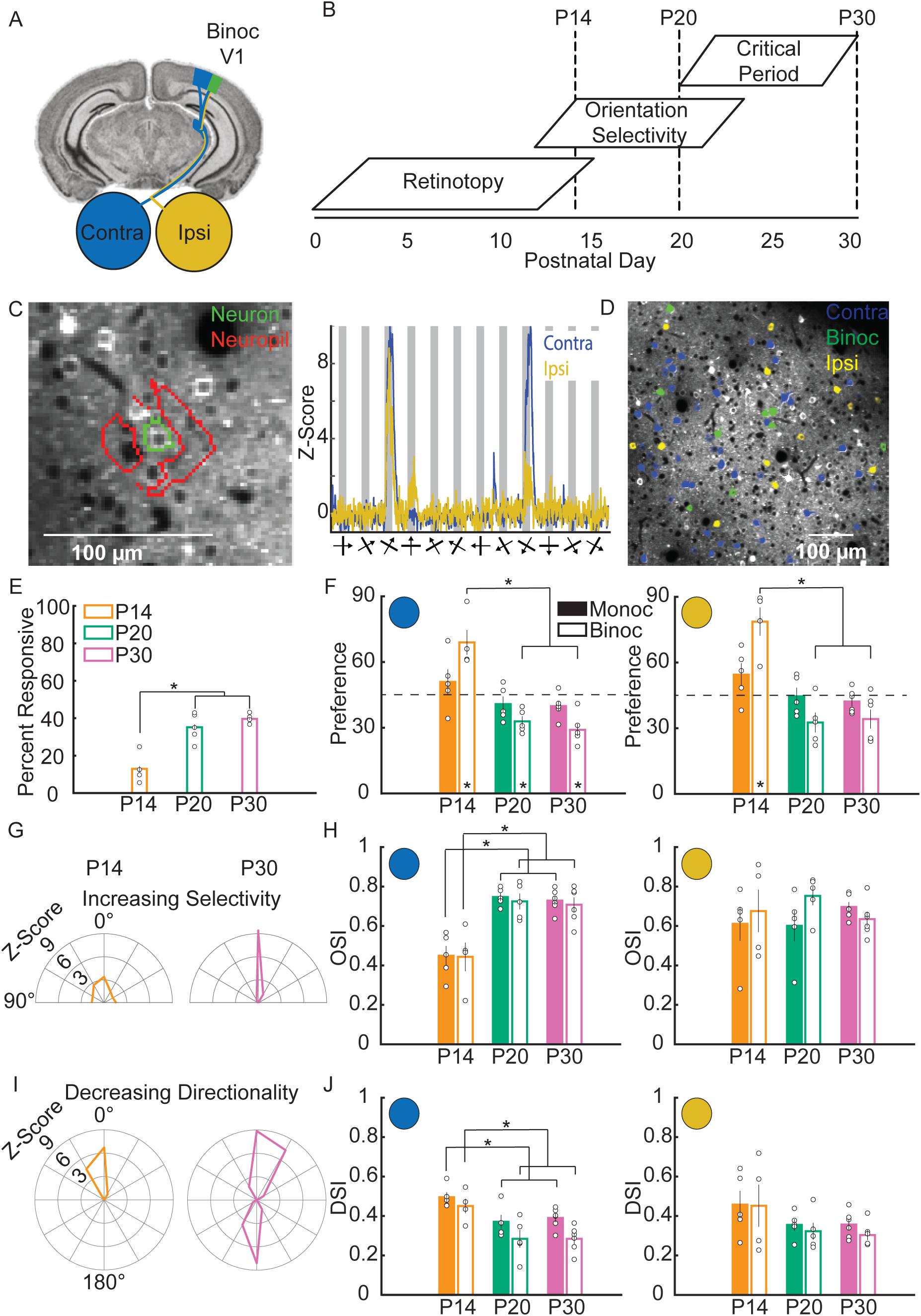
Visual response properties in layer 2/3 excitatory neurons of binocular V1 are rapidly tuned after eye-opening. **A.** Imaging is performed in mouse binocular V1 (green), which receives input from both the contralateral (blue) and ipsilateral (yellow) eyes. **B.** Putative timeline of the development of visual response properties in the mouse. Imaging took place at P14, 20 and 30. **C.** (Left panel) Regions of interest (ROIs) corresponding to the soma of a neuron (green outline) and the surrounding area of neuropil (red outline). (Right panel) The average response of an example neuron, after neuropil subtraction and z-scoring. The thick bars below the X-axis show the orientation of each stimulus while the arrow bisecting it indicates the direction of movement. **D.** Representative image of layer 2/3 excitatory neurons during the recording of visually evoked calcium responses. Neurons that respond solely to the contralateral eye (contra) are labeled blue, neurons that respond solely to the ipsilateral eye (ipsi) are labeled yellow, and neurons that respond to both eyes (binoc) are labeled green. **E.** Percentage of neurons that are visually responsive at each time point. This fraction increases significantly from P14 to later ages. (**F**,**H**,**J**) for all response properties, the left most graphs shows responses of contra-eye stimulation. Monocular neuron responses (solid bars) and binocular neuron responses are shown (empty bars). The right most graphs show the response of ipsi-eye stimulation. **F.** Quantification of orientation preference. Monocular neurons do not change preference over age, while binocular neurons show a shift in orientation preference towards horizontal (0°) from P14 to P20 and P30. Asterisk within a bar denotes an orientation bias away from the expected, unbiased mean preference of 45° (dashed line). **G.** Example Z-scored responses of single neurons to orientations of grating from 0-180° from a P14 and a P30 mouse. Neurons at P30 on average respond more specifically to their preferred orientation (centered on 0°, vertical) than other orientations. At P14, more neurons respond non-specifically to many orientations. **H.** Quantification of orientation selectivity index (OSI). Contralateral responses of monocular and binocular neurons show increases in OSI over age. Ipsilateral responses do not change. **I.** Example Z-scored responses of single neurons to moving grating from 0-360°. Neurons at P14 are on average more direction selective than at P30. **J.** Quantification of direction selectivity index (DSI). Contralateral responses of monocular and binocular neurons decrease in DSI over age. Ipsilateral responses do not change. (* indicates significant difference, p<0.05. Error bars represent standard error of the mean. Open circles indicate a data point from an individual mouse).

### Distinct development of eye specific response properties in monocular and binocular neurons

Little is known about how binocularity is encoded and tuned during development. To determine how visual response properties develop in monocular and binocular neurons, we quantified orientation preference, orientation selectivity, and direction selectivity across age (Figure 1F-J) as these properties are important for visual feature detection. The binocular neuron responses were quantified for each eye. Orientation preference was examined from 0° (horizontal) to 90° (vertical). Responses in binocular neurons, but not monocular neurons, showed a significant change in mean orientation preference towards horizontal orientations from P14 to P20 and P30 (Figure 1F, contralateral monocular; P14=50.9±5.7°, P20=40.8±3.5°, P30=39.9±2.2°, F_2,13_=2.5, p=0.1247, ANOVA. contralateral binocular; P14=69.0±6.3°, P20=33.0±2.3°, P30=29.-±2.7°, F_2,12_=31.8, p<0.0001, ANOVA. Ipsilateral monocular; P14=54.5±5.2, P20=44.6±3.9°, P30=42.2±2.2°, F_2,13_=2.9, p=0.0887, ANOVA. Ipsilateral binocular; P14=78.6±7.2°, P20=32.6±4.5°, P30=34.2±4.3°, F_2,12_=22.5, p<0.0001, ANOVA). At P14, the contralateral and ipsilateral orientation preference of binocular neurons was biased towards vertical orientations (contralateral: p=0.0326. Ipsilateral: p=0.0184). At P20 and P30, binocular neuron contralateral and ipsilateral preference are biased towards horizontal orientations (contralateral: P20 p=0.0066, P30 p=0.0020. ipsilateral: P20 p=0.0523, P30 p=0.0548. *t*-test) while monocular neurons did not display a bias at any age. This suggests that binocular cells develop unique orientation preference as compared to monocular cells.

We next measured orientation selectivity for neurons, quantified as an orientation selectivity index (OSI) (see methods). Both monocular and binocular neuron OSI to the contralateral eye increased significantly from P14 to P20 and P30 (Figure 1H, contralateral monocular; P14=0.45±0.05, P20=0.75±0.02, P30=0.73±0.02, F_2,13_=25.1, p<0.0001, ANOVA. Contralateral binocular; P14=0.44±0.08, P20=0.72±0.04, P30=0.71±0.04, F_2,12_=8.6, p=0.0047, ANOVA). In contrast, OSI to the ipsilateral eye did not significantly change with age in monocular and binocular neurons (Ipsilateral monocular; P14=0.61±0.09, P20=0.60±0.08, P30=0.69±0.02, F_2,13_=0.7, p=0.5154, ANOVA. Ipsilateral binocular; P14=0.68±0.12, P20=0.75±0.05, P30=0.63±0.04, F_2,12_=0.9, p=0.4390, ANOVA). This suggests that OSI tuning develops specifically in contralateral eye inputs. The direction selectivity index (DSI) was calculated for all neurons (see methods). Both monocular and binocular neuron DSI to the contralateral eye decreased significantly over age (Figure 1J, contralateral monocular; P14=0.50±0.03, P20=0.37±0.03, P30=0.39±0.02, F_2,13_=5.9, p<0.0152, ANOVA. Contralateral binocular; P14=0.45±0.04, P20=0.28±0.04, P30=0.28±0.03, F_2,12_=6.0, p=0.0157, ANOVA), while DSI to the ipsilateral eye were unchanged (Ipsilateral monocular; P14=0.46±0.07, P20=0.35±0.03, P30=0.36±0.03, F_2,13_=1.7, p=0.2253, ANOVA. Ipsilateral binocular; P14=0.45±0.12, P20=0.32±0.04, P30=0.30±0.02, F_2,12_=1.5, p=0.2534, ANOVA). Thus, similar to OSI, developmental tuning of DSI occurs specifically in contralateral inputs.

These results reflect complex developmental fine-tuning of visual response properties in binocular V1. Interestingly, the tuning of visual responses did not change significantly from P20 to P30, suggesting many aspects of visual development in binocular V1 mature prior to the start of the classical OD critical period. Tuning of visual response properties occurs mostly through contralateral eye input and binocular neurons become uniquely tuned to horizontal orientations, indicating that these neurons may mediate different processing of visual information.

### The development of binocular neurons requires visual experience

Our results suggest that binocular neurons have specific visual response properties, therefore we investigated these neurons in more detail. Only a small percentage of visually responsive cells were binocular early at P14, with most cells responding only to the contralateral or ipsilateral eye (Figure 2A and B). However, the percentage of binocular cells significantly increased with age (Figure 2B, P14=8±3%, P20=25±3%, P30=16±2%, F_2,13_=9.0, p=0.0036, ANOVA). This was not due to more neurons becoming visually responsive, as the probability of a visually responsive neuron being binocular by chance does not change over age (Figure S1A, P14=0.20±0.03, P20=0.22±0.01, P30=0.22±0.01, F_2,13_=0.6, p=0.5456, ANOVA). We also assessed the relative strength of contralateral and ipsilateral inputs to binocular neurons. The ocular dominance index (ODI) of binocular neurons (see methods) did not change over age (Figure 2C and S1C, P14=0.39±0.08, P20=0.45±0.04, P30=0.38±0.03, F_2,12_=0.5, p=0.6229, ANOVA). This suggests that binocularity at eye opening in mouse V1 is represented at the population level mostly by monocularly driven cells, while individual cells become binocularly responsive later in development.

**Figure 2:**
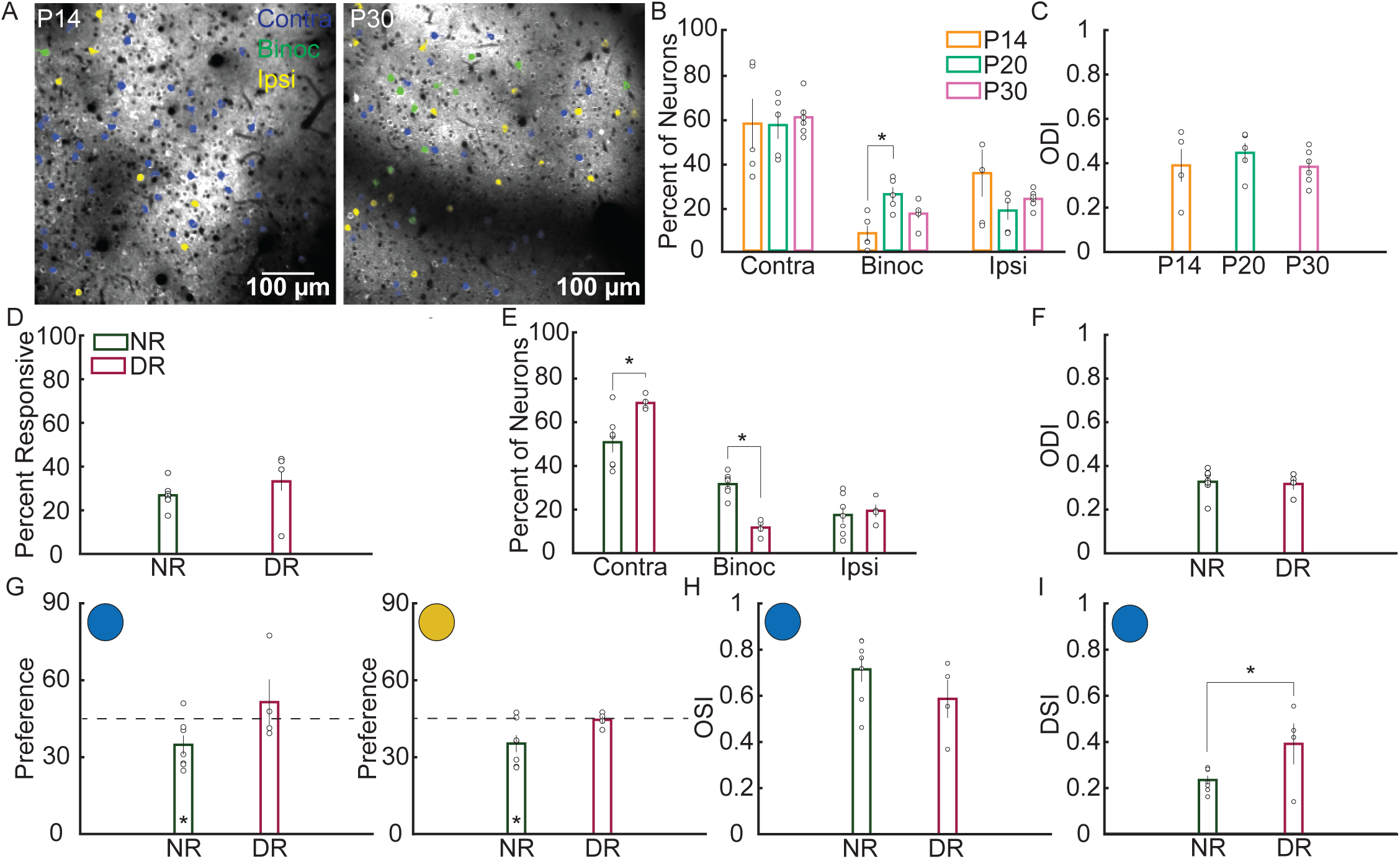
Experience-dependent development of binocular neurons. **A.** Representative images from two-photon recording of visually evoked calcium responses from a P14 (left) and a P30 (right) wild type mouse. **B.** Percentage of visually responsive neurons that respond only to the contralateral eye (contra), to the contralateral and ipsilateral eye (binoc), or only to the ipsilateral eye (ipsi). The percentage of binocular responding cells increases over age. **C.** Ocular dominance index (ODI) of binocular neurons across development. There is no significant difference in ODI at any age. **D.** Percent of neurons responsive to visual stimulation in normally reared (NR) and dark reared (DR) mice. The percent of visually responsive neurons does not differ between NR and DR mice. **E.** The percentage of contralateral neurons is significantly higher in DR than NR mice. The percentage of binocular neurons is significantly lower in DR than NR mice. **F.** There is no significant difference in ODI between NR and DR mice. **G.** Orientation preference of binocular neuron responses to the contralateral eye (left) and ipsilateral eye (right). NR mice show a significant bias for horizontal orientations in either eye, which is not present in DR mice. **H.** OSI did not change with DR. **I.** DSI of the contralateral response of binocular neurons. DSI was significantly higher in DR mice than in NR mice. (* indicates significant difference, p<0.05. Error bars represent standard error of the mean. Open circles indicate a data point from an individual mouse).

The contribution of visual experience to the maturation of visual responses in V1 remains unclear. We therefore investigated whether the emergence of binocular neurons requires visual experience. We dark reared thy1-GCaMP6s WT mice from birth in complete darkness (DR) until P30, with controls raised on a normal 12:12 h light dark cycle (NR). On the day of surgery and recording, all mice were transported, anesthetized, and allowed to recover in darkness until immediately prior to recording. NR and DR mice did not differ in the total percentage of neurons that were visually responsive (Figure 2D, NR=41±4 neurons/mouse N=7, 33.2±8.4%; DR=67±18 neurons/mouse N=4, 26.5±2.2%, F_1,9_=1.0, p=0.3463, ANOVA). We examined the distribution of binocular neurons within the population of responsive neurons (Figure 2E). At P30, DR mice had significantly more monocular contralateral neurons (Figure 2F, NR=50.8±4.6%, DR=68.8±1.7%, F_1,9_=8.1, p=0.0191, ANOVA) and significantly fewer binocular neurons (NR=31.7±1.9%, DR=11.7±1.9%, F_1,9_=46.1, p<0.0001, ANOVA) than NR mice, reminiscent of P14 NR animals (Figure 2B). The ODI of binocular neurons was not altered between NR and DR mice (Figure 2F, NR=0.31±0.02, DR=0.31±0.03, F_1,9_=0.1, p=0.7977, ANOVA). These findings indicate visual experience is crucial to the emergence of binocular neurons.

Similar to the results obtained above with WT mice, this cohort of NR WT mice developed a significant bias towards horizontal orientations in binocular neurons for both contralateral and ipsilateral responses (Figure 2G, Contralateral; 34.8±3.7°, p=0.0327, *t*-test. Ipsilateral; 35.3±3.3°, p=0.0262, *t*-test). In contrast, this bias was not present in DR mice (Contralateral; 51.5±8.8°, p=0.5161, *t*-test. Ipsilateral; 44.5±1.5°, p=0.7775, *t*-test). The contralateral OSI of binocular neurons was not significantly different between NR and DR mice (Figure 2H, NR=0.71±0.05, DR=0.59±0.08, F_1,9_=1.8, p=0.2074, ANOVA). The mean contralateral DSI of binocular neurons, however, was significantly lower in NR mice than DR mice, indicating the developmental decrease in DSI is experience dependent (Figure 2I, NR=0.23±0.02, DR=0.39±0.09, F_1,9_=5.2, p=0.0485, ANOVA). These results suggest that, except for orientation selectivity, fine-tuning of visual response properties in binocular neurons requires visual experience.

### Visual experience drives correlation of binocular neuropil

The neuropil is comprised of intracortical axons and dendrites from L2/3, L4 and L5, as well as thalamic projections into V1, and is densely labeled by GCaMP6s^15^. Since LFP/visually evoked potentials in binocular V1 are predominately binocular, we predicted that neuropil responses may also exhibit binocular responses, despite most single neurons being monocular. However, it is not clear whether neuropil responses would show any developmental-dependent tuning in visual responses or binocularity. Small (7-15 µm radius) neuropil patches in L2/3 of visual cortex have distinct stimulus specific preferences and perform as well as neurons in encoding stimulus direction in adult mice^19^. We examined visual responses of neuropil patches (64-170 µm^2^), taken as an annulus around the soma of all detected neurons 3 times the size of the neuron’s radius (see methods), and found that many patches displayed significant visual responses evoked by either eye (Figure 3A). Similar to neurons, the percentage of neuropil patches with significant visual responses increased over age (Figure 3B, P14=66±11.7%, P20=97.7±0.9%, P30=95.5±2.2%, F_2,13_=7.2, p=0.0079, ANOVA). Given the slow emergence of binocularity at the neuronal level after birth, we predicted that emergence of correlated binocular tuning in the neuropil would also emerge after eye opening. To test this, we measured signal correlation between mean contralateral and ipsilateral responses in binocular neuropil^17,20^. Correlation of the contralateral and ipsilateral responses was low at P14 and P20, but increased significantly at P30 (Figure 3C, P14=-0.08±0.1, P20=0.09±0.07, P30=0.33±0.06, F_2,13_=7.8, p=0.0060). We next examined whether the increase in binocular neuropil correlated activity over development is experience-dependent. Correlation between contralateral and ipsilateral responses in neuropil patches was 0.38±0.07 in NR mice, but completely absent in in DR mice at 0.03±0.03 (Figure 3D, F_1,9_=14.0, p=0.0046).

**Figure 3:**
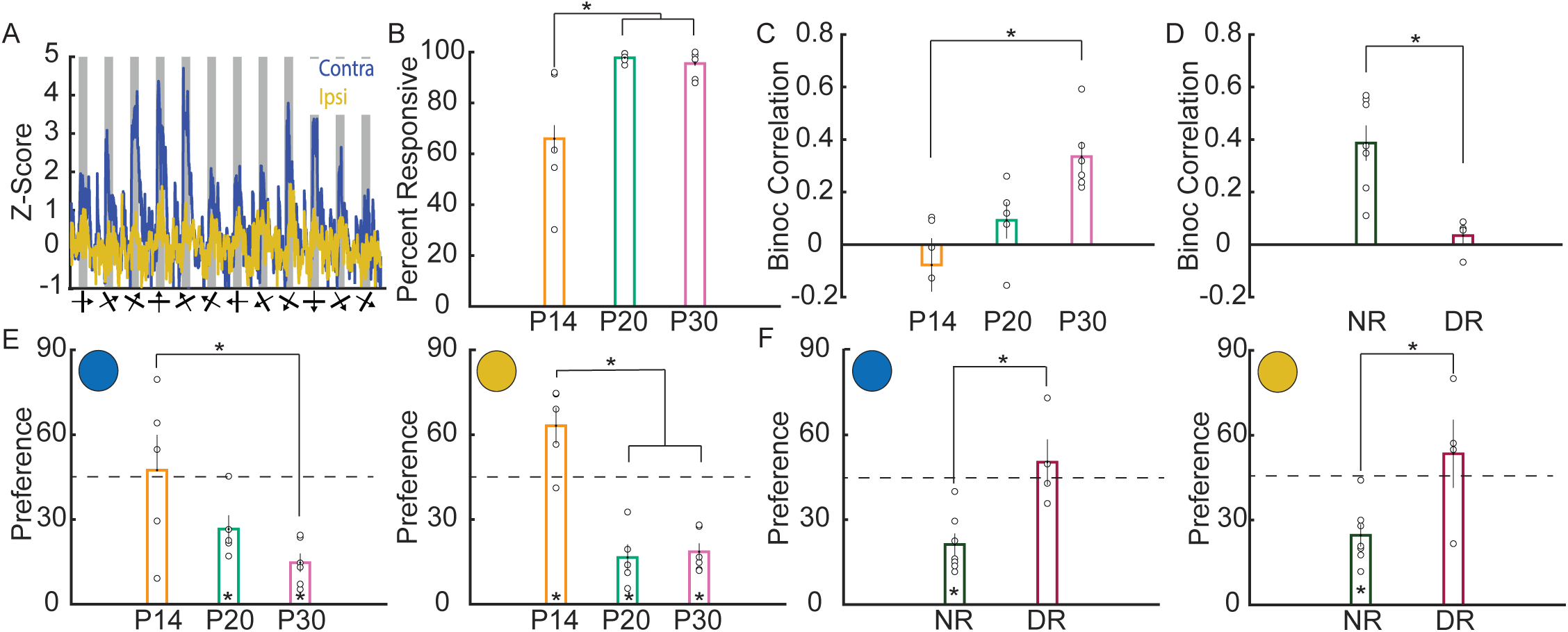
Experience-dependent increases in correlated binocular responses occur in neuropil during development. **A.** Example calcium responses of a binocular region of neuropil to the contralateral (blue trace) and ipsilateral (yellow trace) eye. **B.** Percentage of neuropil surrounding neurons that respond to visual stimuli increases over age. **C.** Signal correlation between the contralateral and ipsilateral responsiveness of binocular neuropil patches increases over age. **D.** The increase in binocular signal correlation is absent in DR mice. **E.** Orientation preference for contralateral and ipsilateral responses of binocular neuropil become biased towards horizontal orientations over age. **F.** Orientation preference is not biased in DR mice. (* indicates significant difference, p<0.05. Error bars represent standard error of the mean. Open circles indicate a data point from an individual mouse).

We next examined orientation preference of neuropil patches across development to determine whether neuropil preference showed similar biases to binocular neurons. Indeed, for both contralateral and ipsilateral responses, mean orientation preference decreased towards 0° from P14 to P30 (Figure 3E, contralateral, left; P14=47.4±12.6°, P20=26.6±4.9°, P30=14.7±3.3°, F_2,13_=4.84, p=0.0269. Ipsilateral, right; P14=63.2±6.4°, P20=16.6±4.6°, P30=18.6±10.8°, F_2,13_=30.6, p<0.0001, ANOVA), and a horizontal bias is apparent at P20 (contralateral: p=0.0200. ipsilateral: p=0.0035, *t*-test) and P30 (contralateral: p=0.0003. ipsilateral: p=0.0003, *t*-test). These results suggest that axons and dendrites within local patches of binocular neuropil develop correlated tuning to visual input from both eyes in a timescale similar to the emergence of single-neuronal binocularity. Additionally, the orientation preference tuning towards horizontal stimuli in neuropil matches that of binocular neurons. Moreover, NR mice showed a horizontal bias for both contralateral (Figure 3F, left, 21.2±3.9°, p=0.0009, *t*-test) and ipsilateral (Figure 3F, right, 24.6±4.0°, p=0.0022, *t*-test) orientation preference, while DR mice showed no bias (contralateral; 50.3±8.1°, p=0.5562. ipsilateral; 53.4±12.0°, p=0.5333, *t*-test). Both contralateral and ipsilateral responses were significantly different from NR mice (contralateral; F_1,9_=13.7, p=0.0049, ANOVA. Ipsilateral; F_1,9_=7.93, p=0.0202, ANOVA). Thus, binocular correlated activity and development of orientation preference in neuropil requires visual experience.

### Arc limits the development of binocular neurons

To gain molecular insight into the processes that regulate the development of binocularity, we investigated the role of the activity-dependent gene *Arc*. We measured visual responses in Arc KO mice at P14, P20 and P30 (Figure 4A). Since this Arc KO line was created by knock in of a destabilized GFP^14^, we examined the fluorescence during grey screen presentations to determine if the GFP signal would lead to higher baseline fluorescence (see methods), possibly biasing analysis of changes in fluorescence. However, the baseline fluorescence in Arc KO mice did not significantly differ from the WT P14, P20 and P30 data (Figure S2, KO P14=34±6 neurons/mouse N=6; P20=47±14 neurons/mouse N=3; P30=59±8 neurons/mouse N=4. P14; WT=214±30, KO=203±17, p=0.7603, *t*-test. P20; WT=315±62, KO=244±3, p=0.3203, *t*-test. P30; WT=278±33, KO=231±6, p=0.2432, *t*-test). The percent of visually responsive neurons responsive solely to the contralateral eye decreased significantly over development from 85.9±1.2% at P14 to 41.5±3.2% at P30 (Figure 4B, P20=70.9±4.2%, F_2,10_=126.9, p<0.0001, ANOVA), whereas binocular responsive neurons increased significantly from 2.0±1.0% at P14 to 29.2±3.9% at P30 (F_2,10_=32.9, P20=11.5±4.4%, p<0.0001, ANOVA). Monocular ipsilateral responsive neurons also increased significantly from 12.0±1.9% at P14 to 29.3±2.3% at P30 (Figure 4B; P20=17.6±1.5, F_2,10_=19.7, p=0.0003, ANOVA). ODI of binocular neurons in Arc KO mice did not significantly change over development (Figure 4C, P14=0.33±0.07, P20=0.38±0.03, P30=0.41±0.02, F_2,7_=0.8, p=0.4719, ANOVA). No difference in the percentage of visual responsive neurons were observed between KO and WT mice at P30 (Figure 4D, WT=39.6±0.9%, KO=41.1±2.8%, F_1,8_=0.3, p=0.5825, ANOVA). Direct comparison of WT and Arc KO mice showed that the percent of visually responsive neurons that were binocular was significantly higher in Arc KO mice than in WT at P30 (Figure 4E, WT=16.6±2.1%, KO=29.2±3.9%, F_1,8_=9.8, p=0.0141, ANOVA), In addition, the percent of visually responsive neurons responding solely to the contralateral eye was significantly lower in Arc KO mice compared to WT mice (WT=60.0±3.5%, KO=41.5±3.2%, F_1,8_=13.7, p=0.0060, ANOVA). ODI did not significantly differ between the two genotypes (Figure 4F, WT=38.4±3.2%, KO=40.6±2.1, F_1,8_=0.3, p=0.6174, ANOVA). We next compared visual response properties of binocular neurons in P30 WT and Arc KO mice. P30 Arc KO mice orientation preference for the contralateral and ipsilateral response of binocular neurons trended towards a horizontal bias (Figure 4G, Contralateral; WT=29.- ±2.7°, KO=29.8±4.8°, p=0.0516. Ipsilateral; WT=34.2±4.3°, KO=30.1±5.3°, p=0.0684, *t*-test), and did not significantly differ from WT (Contralateral: F_1,8_=0.02, p=0.8825, ANOVA. Ipsilateral: F_1,8_=0.36, p=0.5670, ANOVA). Similarly, neither contralateral OSI (Figure 4H, WT=0.71±0.04, KO=0.71±0.05, F_1,8_=0.0, p=0.9987, ANOVA) or contralateral DSI (Figure 4I, WT=0.28±0.03, KO=0.23±0.01, F_1,8_=2.7, p=0.1384) differed between P30 WT and KO.

**Figure 4:**
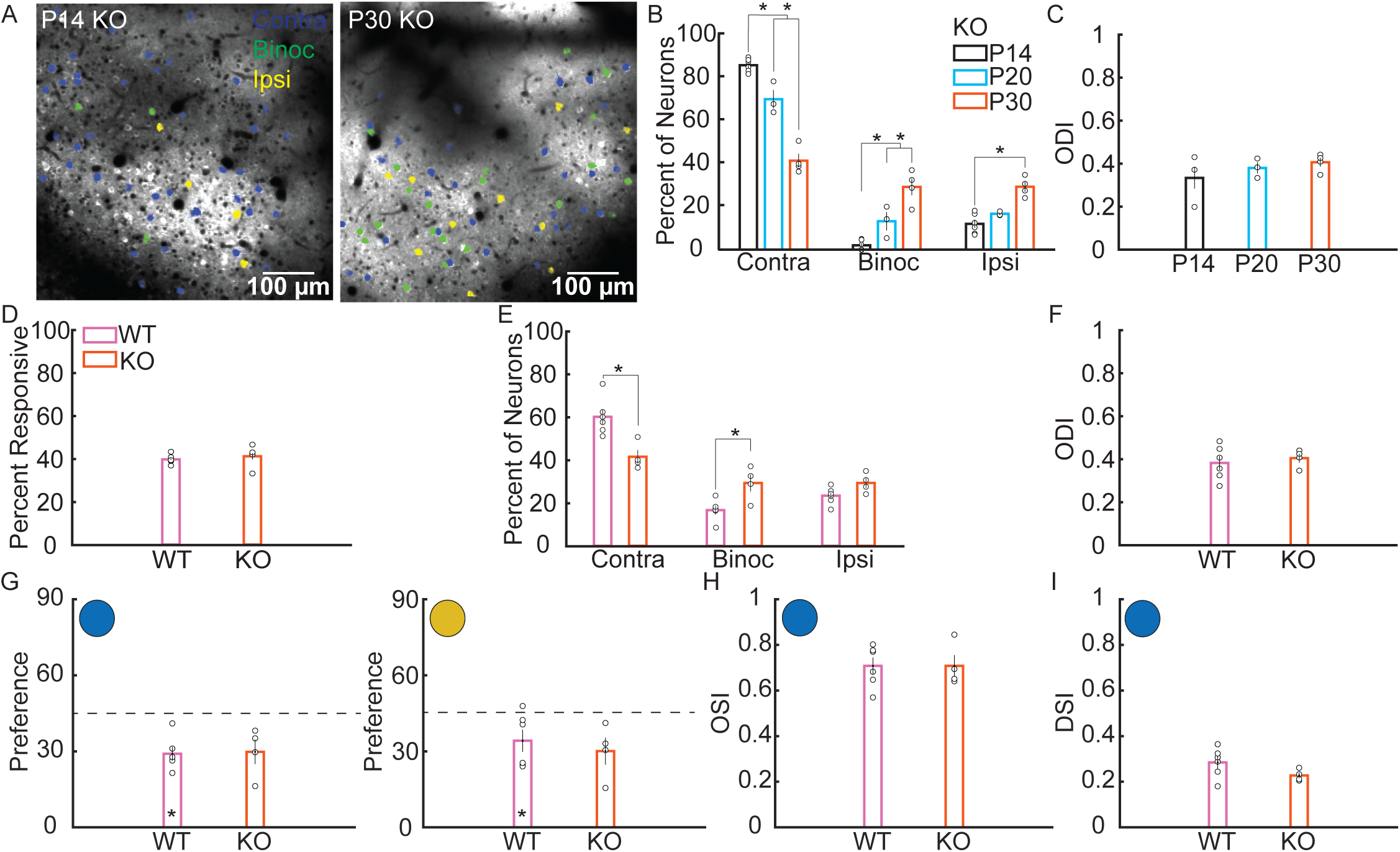
Arc limits the emergence of binocular neurons early in development. **A.** Representative images from two-photon recording of visually evoked calcium responses in Arc knock out (KO) mice. **B.** Percent of visually responsive neurons that respond only to the contralateral eye (contra), to the contralateral and ipsilateral eye (binoc), or only to the ipsilateral eye (ipsi) in Arc KO mice. The percent of contra neurons decreases from P14 to P20, and from P20 to P30. The percent of binoc neurons increases significantly from P14 to P20, and from P20 to P30. The percent of ipsi neurons increases significantly from P14 to P30. **C.** ODI of binocular neurons does not change across development in Arc KO mice. **D.** Comparison of littermate P30 WT and Arc KO mice. The percent of visually responsive neurons did not significantly differ between P30 WT and Arc KO mice. **E.** Percent of visually responsive neurons that are contra, binoc, or ipsi responsive in P30 WT and Arc KO mice. The percent of contra neurons is lower in Arc KO mice, while the percent of binoc neurons is higher. **F.** There is no significant difference in ODI of binocular neurons between WT and Arc KO mice. **G.** Both WT and Arc KO mice show a bias for horizontal orientations in either eye. **H.** OSI of contralateral responses did not significantly differ between P30 WT and Arc KO mice. **I.** DSI of contralateral responses was not significantly different between P30 WT and Arc KO mice. (* indicates significant difference, p<0.05. Error bars represent standard error of the mean. Open circles indicate a data point from an individual mouse).

Surprisingly, rather than mirroring the effect of dark rearing, loss of Arc results in an increase in binocular neurons without impacting other response properties. This suggests that Arc may tune cortical circuits during developing by acting as a break on eye-specific synaptic plasticity.

### Binocularity in the adult visual cortex requires ongoing maintenance by Arc-dependent synaptic plasticity

Arc protein expression peaks during the critical period^13^, thus we predicted that Arc’s role in regulating binocular neurons would be constrained to early development. However, OD plasticity still occurs in adult mouse visual cortex, although primarily through potentiation of open eye/ipsilateral responses^21^, suggesting the balance of input between the two eyes is still plastic and may require ongoing maintenance. To determine whether Arc is required for the maintenance of binocular cells in adult binocular V1, we used a floxed Arc line (Arc cKO) mice^22^, that was crossed into the thy1-GCaMP6s line. At P180 we injected either AAV5-hSyn-mCherry (Control) or AAV5-hSyn-mCherry-Cre (Cre) virus into L2/3 of binocular V1. Two weeks following injection, we performed a craniotomy over the injection site and imaged as performed for young mice. Immunohistochemistry of the injected hemisphere confirmed effective recombination of Arc after mCherry-Cre expression (Figure S3A). As in P30 mice, a mix of monocular and binocular neurons were visible in L2/3 for both control and Cre injected mice (Figure 5A). Strikingly, Cre mediated knockdown of Arc in adult binocular V1 was sufficient to alter the percentage of binocular neurons as compared to control injected mice (Figure 5B, Control=55±7 neurons/mouse N=5; Cre=56±6 neurons/mouse N=5; Control=12.6±2.5%, Cre=26.6±5.3%, F_1,8_=5.7, p=0.0442, ANOVA). Additionally, Cre injected mice showed a decrease in the number of monocular contralateral neurons (Control=70.7±4.8%, Cre=40.6±6.2%, F_1,8_=14.7, p=0.0050, ANOVA) and an increase in monocular ipsilateral neurons (Control=16.6±4.1%, Cre=32.8±3.6%, F_1,8_=8.8, p=0.0180, ANOVA). ODI of binocular neurons was unaffected (Figure 5C, Control=40.8±2.1%, Cre=35.2±2.7%, F_1,8_=2.6, p=0.1442, ANOVA). We next compared response properties of binocular neurons in Control and Cre injected mice. Contralateral orientation preference of Control mice trended towards a horizontal bias (Figure 5D, 29.7±5.7°, p=0.0549, *t*-test), and ipsilateral orientation preference was significantly biased towards horizontal orientations (27.6±4.7°, p=0.0213, *t*-test). Cre injected mice had a significant horizontal bias of the binocular neuron’s contralateral response (36.9±2.3°, p=0.0261, *t*-test) and did not differ significantly from Control (F_1,8_=1.4, p=0.2745, ANOVA). However, Cre mice did not show a bias for the ipsilateral response (41.1±2.0°, p=0.1209, *t*-test) and significantly differed from Controls (F_1,8_=7.0, p=0.0295, ANOVA). Neither contralateral OSI (Figure 5E, Control=0.81±0.02, Cre=0.74±0.04, F_1,8_=2.6, p=0.1466, ANOVA) or contralateral DSI (Figure 5F, Control=0.24±0.05, Cre=0.26±0.02, F_1,8_=0.1, p=0.7430) differed between Control and Cre. See supplementary table 1 for a summary of all response properties across genotype and manipulation.

**Figure 5:**
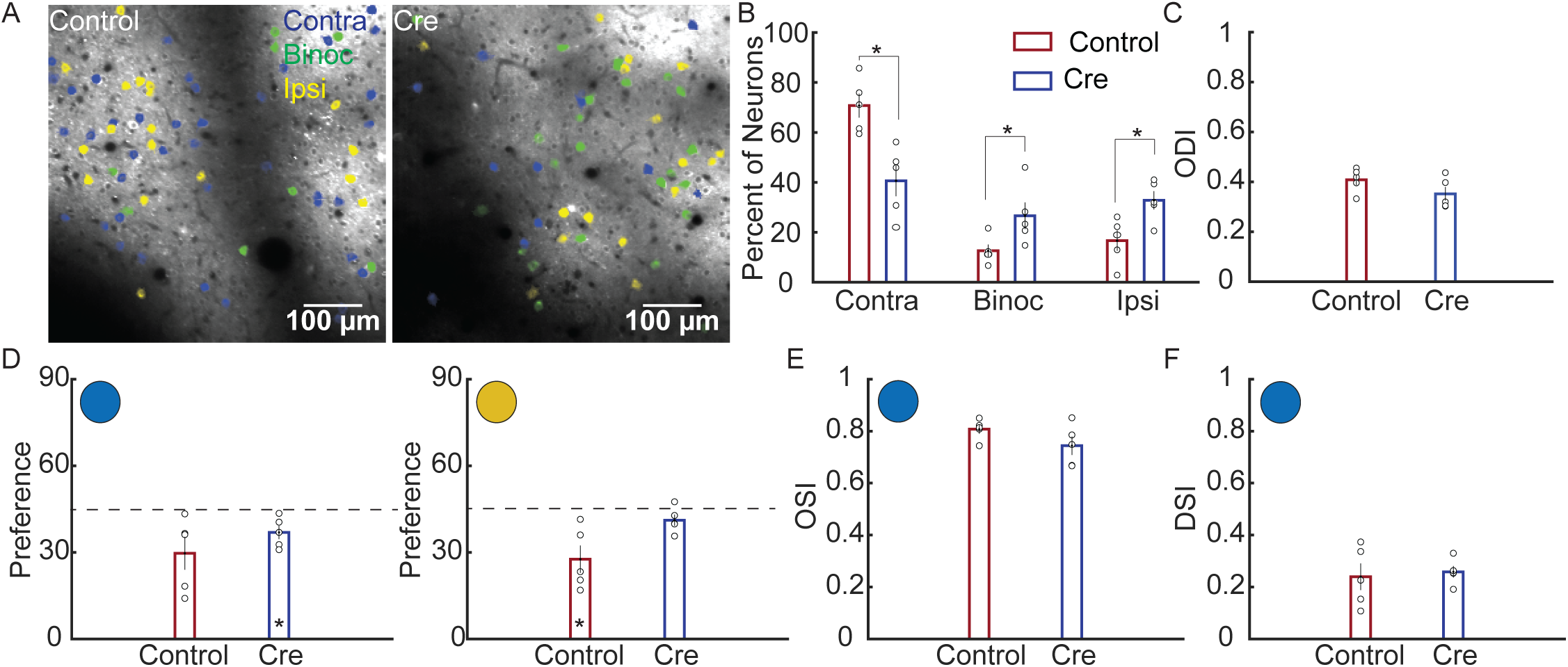
Arc is required for the maintenance of binocular neurons in adult binocular V1. **A.** Representative images from two-photon recording of visually evoked calcium responses from neurons imaged in a Control and Cre injected mouse. **B.** Percent of visually responsive neurons that are contra, binoc or ipsi responsive. The percent of contra neurons is significantly lower in Cre than in Control mice, while the percent of binoc and ipsi neurons is higher. **C.** There is no significant difference in ODI of binocular neurons between Control and Cre mice. **D.** Orientation preference of binocular neuron responses to the contralateral (left) and ipsilateral (right) eye. Cre mice show a small but significant horizontal bias in contralateral responses. Control mice show a horizontal bias in ipsilateral responses. **E.** OSI of contralateral responses did not significantly differ between Control and Cre mice. **F.** DSI of contralateral responses did not significantly differ between Control and Cre mice. (* indicates significant difference, p<0.05. Error bars represent standard error of the mean. Open circles indicate a data point from an individual mouse)

These results suggest that the precise regulation of binocularity in V1 requires ongoing maintenance that is dependent on Arc expression, even in adult mice.

## DISCUSSION

The interplay between experience and genetic hardwiring of circuits in the brain results in optimal processing of information from the outside world. Here, we show that at eye-opening the mouse binocular V1 is mostly driven by monocular responding cells, while binocular cells rapidly emerge after eye opening in an experience-dependent process that is regulated by the activity-dependent gene *Arc*. These binocular neurons convey specific visual response properties, including a preference for horizontal orientations. The increase in binocular responding neurons is mirrored by an increase in the signal correlation of contralateral and ipsilateral input in the neuropil, which is also experience-dependent. These data solidify the role of experience as a crucial component of refining sensory processing and binocular vision early in development.

While it is clear that there are critical windows of plasticity during development that are essential for correct wiring of neuronal circuits, it has also become evident that brain plasticity still occurs in adult brains^23^. Indeed, many studies have shown that juvenile-like plasticity can be reinstated in adult brains^13,24–27^, and that visual deprivation can alter plasticity rules in adult animals^28^. However, few studies have shown a direct role for plasticity in regulating or maintaining circuits required for sensory processing. Unexpectedly, we found that knock-down of Arc in adult mouse V1 alters the distribution of monocular and binocular neurons that recapitulates the developmental phenotype observed in young Arc KO mice. This suggests that sensory circuits may require ongoing plasticity that refines information processing in response to experience.

### Binocularity develops after eye opening

Most studies that have evaluated binocular mouse visual cortex have used methods that lack single-cell resolution (intrinsic imaging, local field potentials) or have issues with sampling bias (single unit electrophysiology). We find, at the single cell level, that binocular visual cortex is mostly comprised of monocular responding cells (∼90%) initially at eye-opening and that by P30 there are ∼80% monocular/∼20% binocular cells, which remains constant into adulthood. This high prevalence of monocular cells in binocular visual cortex is similar to what has been observed using GCaMP6 recordings^7,29^. However, it is possible that because transgenic thy1-GCaMP6 expression is lower than viral expression of GCaMP6 we may miss very weak responses, especially from the ipsilateral eye that would lead to under-estimation of binocular cell numbers^15^.

While the percentage of binocular single-cell responses is low, the emergence of binocular cells is correlated with an increase in tuning between contralateral and ipsilateral input in the neuropil. Calcium signals in large (∼10,000 µm^2^) sections of neuropil correlate well with electro-corticogram measures of sensory evoked population activity^30^. However, smaller sections of neuropil can report stimulus specific information that have response properties comparable to neurons in the same area^19^. While this local specificity is incompatible with the view of mouse V1 as a “salt and pepper” mix of orientation preference, this finding agrees with the observation that neurons with similar tuning are arranged in mini-columns and that thalamacortical projections from LGN to V1 are arranged in patches rather than having no spatial organization^8,31^. Therefore, the axons and dendrites making up local patches of neuropil likely respond to similar stimuli despite the fact that they arise from different cell bodies. Similarly tuned neurons also preferentially wire together, and the probability for similarly tuned neurons to be connected doubles over development from P13-15 to P22-26^20^. Our findings demonstrate that spatial organization of orientation preference becomes “aligned” between the two eyes over the course of development, mirroring the emergence of binocular neurons. The neuropil activity of the mice imaged in this study are a heterogenous mix of axons and dendrites from different layers and brain regions. It will be of interest to examine the refinement of visual response properties specifically in and projecting to L2/3 of binocular V1 using layer or region specific labeling.

The development of inhibitory neurons plays an important role in regulating V1 plasticity^32^. Parvalbumin positive interneurons make up the majority of cortical interneurons, and from the pre-critical to critical period in binocular V1 display an experience dependent increase in firing rates coinciding with a decrease in orientation selectivity that is thought to be necessary for initiation of the critical period^33^. This reflects integration from the local (∼100 µm) excitatory neuronal population, leading to a strongly binocular, but non-selective inhibitory population^34^. The experience dependent emergence of excitatory binocular neurons we observe may be what initiates this net binocular drive onto inhibitory neurons.

Since our recordings are acute, we were not able to follow the same neurons over time. Our data suggests that binocular neurons do not arise solely from neurons that have become visually responsive. Thus, neurons could become binocular by the addition or strengthening of ipsilateral eye input into V1, i.e. a switch from monocular contralateral to binocular. Alternatively, neurons that start off as monocular ipsilateral could switch to binocular by the addition or strengthening of contralateral input.

We find that binocular neurons have different response properties to that of monocular neurons. In particular, binocular cells rapidly develop an orientation preference bias towards horizontal stimuli that was not observed in monocular neurons. Interestingly, we found that orientation and direction selectivity changes during development are only observed in contralateral eye responses, both in monocular and binocular neurons. This suggests both eye-specific information and plasticity is required to refine binocular V1. We conclude that the emergence of binocular responding cells is key for processing of input from each eye.

### Experience shapes the development of binocularity

Much of what is known about experience-dependent plasticity in V1 comes from monocular deprivation studies. Much less is known about the normal activity-dependent processes that are required for normal visual response properties. In cats, where the majority of neurons in V1 are binocularly responsive, spontaneous activity originating in the retina is sufficient to guide the formation of OD bands^35^. However, visual experience is necessary during the critical period in the cat to strengthen ipsilateral responsiveness^36^. We find that the development and tuning of binocular neurons is experience-dependent. Both the increase in binocular neurons and the horizontal orientation preference in the few existing binocular neurons is absent in dark reared mice. Moreover, the increase in correlated binocular activity in the neuropil is also experience-dependent. Interestingly, orientation selectivity does not seem to be experience-dependent. This may be due to processes that occur prior to eye-opening or that tuning occurs predominantly in layer 4 or even thalamic input from the lateral-geniculate nucleus^29^.

Experience is tightly coupled to the induction of activity-dependent gene expression. The immediate early gene Arc is necessary for experience-dependent synaptic plasticity in mouse visual cortex and Arc expression is sufficient to prolong or re-open the critical period of OD plasticity^13^. However, it was not clear how Arc regulates the normal development of visual response properties. Here, we find that Arc expression is required for the normal distribution of monocular and binocular neurons in V1. Surprisingly, Arc KO mice show a significant increase in binocular neurons as compared with WT mice, which is the opposite of what occurs in DR mice. In addition, Arc KO mice show decreased monocular contralateral neurons. Thus, at the population level there is a change in overall OD where contra- and ipsilateral drive is more equal than the contralateral dominance observed in WT mice. Consistent with this, ODI derived from visually evoked potentials shows that Arc KO mice at the population level have ∼1:1 contra to ipsi response amplitude ratio as opposed to ∼2:1 in WT mice^12^.

Arc has been implicated in various cellular models of synaptic plasticity, such as LTP/LTD and homeostatic scaling^37^. It is possible that the phenotypes observed in Arc KO mice are due to a lack of ipsilateral pruning/depression and/or potentiation/strengthening of contralateral inputs. Arc may also have a systems-wide role in regulating experience-dependent correlated activity in different brain regions during development^38^, which may occur through non cell-autonomous regulation of synaptic plasticity through intercellular spread of Arc protein capsids^39^. Further studies will be required to determine precisely how Arc regulates cellular changes in V1 during development.

### Sensory circuits in visual cortex are maintained by synaptic plasticity in adult animals

Experience-dependent plasticity has been well documented in the adult brain, although in general the extent and ease of induction is limited compared with the juvenile brain^23^. Therefore, the role of experience-dependent plasticity in regulating adult sensory processing is thought to be limited. However, manipulations of experience such as dark exposure can reactivate developmental plasticity and allow recovery from developmental deficits such as amblyopia^40^, suggesting that adult cortex is still primed to undergo experience-dependent plasticity. Here, we find that reducing Arc levels only in adult cortex and well-beyond the classical critical period in mice resulted in a significant change in the distribution of monocular and binocular neurons. These changes were observed in a relatively short time course of Arc knockdown (∼2 weeks) and recapitulate the developmental phenotype observed in Arc KO mice. These results suggest that Arc-dependent, and presumably experience-dependent, plasticity is required for ongoing maintenance of binocular circuits in adult V1. These results also suggest that despite a reduction in the induction of Arc protein expression over age^13^, cortical neurons are acutely sensitive to changes in Arc expression.

Taken together, we conclude that experience is required for the emergence and tuning of binocular neurons in V1 during development and for ongoing maintenance in adult V1.

## ACKNOWLEDGEMENTS

We thank Dr. Richard Palmiter (University of Washington) for sharing the Arc conditional KO mouse line. We thank the Wachowiak lab for helpful discussions and technical help with two-photon imaging. K.R.J. was supported by the University of Utah Neuroscience Training Program (5T32NS076067) and by a NIH National Research Service Award F31 (MH112326). This work was funded by the NIH (R00-NS076364 and R01-MH112766), and the E. Matilda Ziegler Foundation.

## AUTHOR CONTRIBUTIONS

K.R.J performed all experiments, wrote code and analyzed the data. K.R.J and J.D.S conceived, designed, and interpreted the experiments. K.R.J and J.D.S wrote the manuscript.

## METHODS

### Animals

All procedures were performed in compliance with the Institutional Animal Care and Use Committee at the University of Utah. The mouse line C57BL/6J-Tg(Thy1-GCaMP6s)GP4.3Dkim/J (Stock No. 024275, The Jackson Laboratory) was used for dark rearing experiments and crossed to Arc KO and Arc cKO mouse lines in all other experiments. The Arc KO (GFP knock in) mouse line C57BL/6-Arctm1Stl/J (Stock No. 007662, The Jackson Laboratory) was used in developmental experiments, with Arc^WT/GFP^Thy1 GCaMP6s pairs set up to yield littermate WT and Arc KO (Arc^GFP/GFP^) mice. Arc conditional knock out (cKO) mice were generously provided by Dr. Richard Palmiter^22^. All mouse lines were maintained on a C57BL/6 background. Both male and female mice were used. Mice were group housed until the day of recording. P14 and P20 mice were kept with their dams until the day of recording. All P30 and older mice were weaned at P21. All mice, with the exception of dark reared mice, were kept on a 12h:12h light/dark cycle. Dark reared mice were housed in a separate room in a dark cabinet and checked daily using a red LED light.

### Cranial window Surgeries

Cranial window surgeries were performed identically for all ages and genotypes used in the study. On the day of recording, mice were anesthetized with 2% isoflurane and injected subcutaneously with enrofloxacin (VETONE), Carprofren (Zoetis), and Dexamethasone (VETONE). The fur over the scalp was trimmed and mice secured in a stereotax (Kopf). Lidocaine (VETONE) was injected subcutaneously under the scalp and the scalp sterilized with alternating swabs of iodine and 70% ethanol. The skin over the skull surface was removed using scissors, and the skull scraped clean using a scalpel and then dried. A 3 mm circle was drawn over the right binocular visual cortex, centered 3mm lateral of midline and 1 mm anterior lambda. The skull outside the 3mm circle and skin margins were covered by a layer of vetbond (3M) and then dental cement (Lang) mixed with black acrylic paint powder (Sargent Art) and allowed to dry. A 3 mm craniotomy was carefully drilled using a high speed drill and 0.5 mm drill bit. Ice cold ACSF (124 mM NaCl, 3.2 mM KCl, 1.25 mM NaH_2_PO_4_, 2 mM CaCl) was periodically placed on the site of drilling to cool the brain and clean the skull surface. When only a thin layer of bone remained at the margins of the craniotomy, forceps were used to remove the circle of bone under a drop of ACSF. ACSF was used to wash the brain surface until surface bleeding was controlled. In some cases, Gelfoam (Pzfizer) was placed on the brain surface to remove excess blood. When bleeding had stopped, a 3 mm circular coverslip (No. 0 coverglass, Warner Instruments) was placed on the brain surface and held down using a needle attached to the arm of the stereotax. Vetbond and dental cement were used to attach the edges of the coverglass to the skull. A custom stainless steel headplate was attached to the skull over the coverglass using dental cement. Animals were allowed to recover on a heating pad and given a post-surgery injection of carprofen subcutaneously to control pain. All animals were allowed to recover for 3 hours minimum prior to recording. In the case of dark reared and normally reared mice, this recovery took place in a dark cabinet and transportation to and from surgery and to the recording room were carried out in the dark with no light exposure until the animal was under anesthesia.

### Two-photon calcium imaging and visual stimulation

Calcium responses were recorded using a raster-scanning two-photon microscope (Bruker Ultima) controlled by Prairie View software. A Coherent Chameleon laser was used for fluorophore excitation, fixed at 920 nm for GCaMP6s imaging and 1020 nm for mCherry imaging. Laser power was set at 80 mW at the sample. A 20x water immersion objective (Olympus, 1 numerical aperture) was used in all experiments. Images were acquired at 2.64-2.94 Hz over a field of view 594×594 µm at 2.32 µm/pixel. All imaging was performed at a depth of 150-210 µm below the pia as measured from the center of the field of view. A dichroic mirror was used to separate red and green fluorescence. All testing was done in awake, head fixed mice held stationary in a plastic tube. Mice were anesthetized with 2% isoflurane and placed below the objective and allowed to habituate for 30 minutes prior to imaging. Normally reared and dark reared mice did not have their monitor turned on until 5 minutes prior to recording. Visual stimuli were presented on a LCD monitor centered perpendicular to the mouse’s field of view, 36.2 cm from the mouse’s head. Visual stimuli consisted of drifting, sinusoidal grating of spatial frequency 0.05 cycles/degree moving 1 cycle/second generated by the MATLAB (Mathworks) toolbox, Psychtoolbox. 6 orientations from 0-150°, each with two directions of movement orthogonal to their orientation, were used to elicit calcium responses. Each direction was repeated 5 times in pseudorandom order. Visual stimuli were each presented for 5 seconds with 5 seconds of gray screen between each presentation. Visual stimuli were time locked to data acquisition using a 5 volt square wave pulse generated using a Measurement Computing USB-1208FS-Plus data acquisition board and recorded by Prairie View. Visual stimuli were presented independently to the contralateral or ipsilateral eye by placing an opaque eye block in front of the opposing eye to temporarily occlude vision. The eye stimulated first (contralateral or ipsilateral) was randomized between animals and recordings.

### Analysis of two-photon imaging data

Only one FOV was quantified for each animal. Collected frames were opened in ImageJ (NIH) using the Prairie Reader plugin, and contralateral and ipsilateral recordings concatenated. The two-photon calcium imaging analysis pipeline, Suite2P (https://github.com/MouseLand/suite2p) was used to register and detect cells in recorded data. Default settings for the December 2018 distribution of Suite2P were used with the exception of the following: diameter of 9, Tau of 2, frames per second 2.64 or 2.94, 3 neuropil to cell ratio and 191 minimum neuropil pixels. Detected ROIs were manually curated to discard ROIs in which a neuronal somatic ROI was not visible in the mean image. Remaining ROIs were classified as putative neurons active during the recording period. All further analysis was carried out using custom functions written in MATLAB (https://github.com/Shepherdlab/2Photon). For each ROI, the corresponding neuropil signal was subtracted from the ROI signal to remove contamination from out of focus fluorescence. The resulting ROI trace was then normalized to the mean and standard deviation of the signal during the gray screen presentations to produce a Z-score ((F-F_mean_)/F_Standard Deviation_). The period preceding and including the 5 presentations of each unique stimulus was then segmented. The means of each presentation and its corresponding pre-stimulus period were statistically compared using a paired sample *t*-test. The 5 stimulus presentations were then averaged together as a robust mean, and the mean Z-score of the mean trace during the stimulus calculated. Only neurons with a significant response to at least one stimulus (during stimulation of either the contralateral or ipsilateral eye) were included in further analysis. The criteria used to determine a significant response was a p-value of the paired *t*-test below 0.05, and a mean Z-score of the mean response above 0.5 for the same unique stimulus. This amplitude threshold was determined by taking the 99.58^th^percentile of mean responses during grey screen (no stimulus periods) for all ages, giving a 0.42% false discovery rate per stimulus, or 5% for each ROI. A ratio of the number of neurons responding to the contralateral eye and the number of neurons responding to the ipsilateral eye was calculated for each mouse. Within each group (genotype/age), this measure was used to remove outliers more than 1 standard deviation from the mean of the group.

Responsive neurons were classified based on whether there was a significant response to stimulation of both the contralateral and ipsilateral eye (Binoc), or only to one eye (Contra or Ipsi). For all responsive neurons, their direction selectivity was calculated using a global measure of direction selectivity equal to

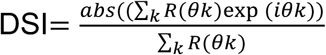

Where R(θk) is the response to the direction θk. Orientation selectivity was calculated in a similar manner, where

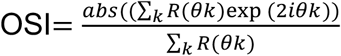

Where R(θk) is the response to the orientation θk. Orientation preference is derived during the above equation as *abs*((∑)R(*θk*)exp (*2iθk*)), and converted to degrees. For analysis using averaged orientation preference of the population, values larger than 90° were subtracted from 180 to align to a 0-90° scale. The ocular dominance index (ODI) of binocular neurons, taken as a ratio of their ipsilateral response over the sum of their ipsilateral and contralateral response, utilized the mean response to the direction which elicited the largest mean response for each eye. To compare the amplitude of responses between groups, ΔF/F_mean_ was used with a neuropil correction factor of 0.7^15^. This was necessary as a Z-scoring does not allow quantitative comparison of response amplitude between animals. Due to the resting fluorescence of Thy1-GCaMP6s being at or below background fluorescence for most cells, Z-scoring was deemed most appropriate to quantify response properties within animals. All data are first averaged within animal and then between animals in a group (age/genotype) unless otherwise noted. All response properties calculated for groups used in this study are reported in supplementary table 1.

For analysis of neuropil patches, in each recording a blood vessel passing perpendicular to the plane of imaging was manually selected and the mean fluorescent signal of the blood vessel in each frame calculated and subtracted from the trace of each neuropil patch. This subtraction controls for any light contamination related to the visual stimuli, and assures neuropil signal reflects true, local calcium fluctuations. Calculation of Z-score and defining significant visual responses were done identically for neuropil and neurons. Signal correlation between the contralateral and ipsilateral responses were calculated as a linear Pearson correlation between vectors of the mean response to each direction in MATLAB^17,20^.

### Virus injections

For Arc conditional knock out studies, either rAAV5/hSyn-mCherry or rAAV5/hSyn-mCherry-Cre were injected at concentrations of 10^12^GC/mL (University of North Carolina Viral Vector Core). Imaging in Thy1 GCaMP6s Arc cKO mice was performed two weeks after virus injection. Mice were anesthetized with 2% isoflurane (VETONE) and injected subcutaneously with enrofloxacin (7mg/kg, VETONE), Carprofren (5mg/kg, Zoetis) and Dexamethasone (0.2 mg/kg, VETONE). The fur over the scalp was trimmed and mice secured in a stereotax (Kopf). Lidocaine (VETONE) was injected subcutaneously under the scalp and the scalp sterilized with alternating swabs of iodine and ethanol. A small incision was then made to expose the skull surface and the position of the right, binocular visual cortex marked (3 mm lateral midline, 1 mm anterior Lambda). A high speed drill (Foredom) with a 0.5 mm drill bit was used to drill a burr hole over the marked site. Virus was loaded into a pulled, glass pipette fitted to a Nanoject II system (Drummond Scientific). The pipette tip was lowered 200 µm from the brain surface and allowed to rest for 5 minutes. 9.2 nLs of virus was injected every 15 seconds for 25 minutes for a total volume of 920 nLs. Following injection, the pipette tip remained in position for 5 minutes before carefully being withdrawn. The injection site was then sealed with a small drop of Vetbond (3M), and the scalp sewn up using vicryl sutures (Ethicon). Animals recovered on a warm heating pad prior to being returned to their cage. Carprofen tablets (2 mg/tablet) were used post-operatively to control pain. Injected mice were returned to their cage with gender matched siblings following surgery.

### Immunohistochemistry

To confirm effective knock out of Arc using mCherry-Cre virus, a P20 Arc cKO mouse was injected at P20 and sacrificed at P34, then perfused with PBS followed by 4% ice cold paraformaldehyde in PBS. The brain was removed and stored in 4% paraformaldehyde in PBS for a minimum of 24h before being transferred to 30% sucrose in PBS. The brain was cut into 40 µm sections on a Leica 1950 CM cryostat and stored in PBS at 4°C. Sections containing binocular visual cortex were blocked in 5% normal donkey serum (Jackson ImmunoResearch)/0.1% Triton X-100 (Amresco) in PBS for 1 hour and then transferred into 1:1000 custom made rabbit anti-Arc antibody (Proteintech) diluted in blocking buffer overnight. Sections were then washed 3 times for 10 minutes each time in PBS, before being placed in secondary donkey anti-rabbit Alexa Fluor 488 (Jackson ImmunoResearch) for 4 hours. Sections were then washed 3 times for 10 minutes each time in PBS before mounting on slides with ProlongGold Antifade reagent (Invitrogen).

### Statistics

N in all analyses is number of animals unless otherwise noted. Calculation of paired *t*-test between pre and post stimulation means on individual trials was calculated using built in MATLAB functions. All other statistics were calculated using JMP (SAS). Significance level was set at α<0.05 for all one/two group comparisons using a two sample ANOVA or *t*-Test. Otherwise, an ANOVA was used to calculate a F value and p value followed by a post-hoc Tukey’s Honest Significant Difference test to determine significant differences between groups.

## SUPPLEMENTARY MATERIAL

**Supplementary Figure 1 related to Figure 1.**
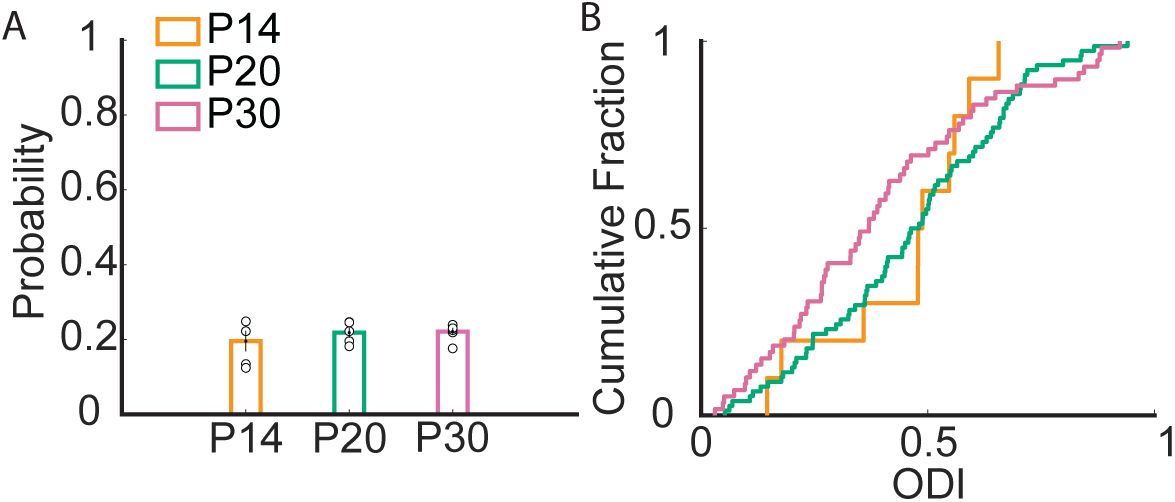
Development of visual response properties. **A.** Theoretical probability of a visually responsive neuron being binocular given the number of contralaterally and ipsilaterally responsive neurons at each age. The mean probability does not change over development. **B.** Cumulative fraction of ODI for all binocular neurons at each age (P14, P20 and P30). There is no significant difference at any age (K.S test).

**Supplementary Figure 2 related to Figure 4.**
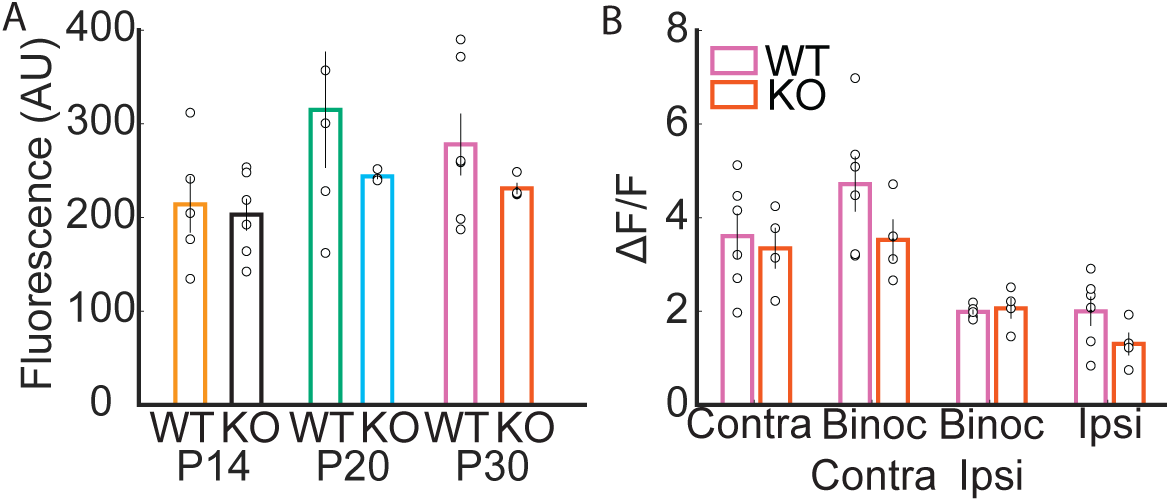
GFP expression in Arc KO neurons does not impact visual response amplitude. **A.** The lowest 5^th^ percentile of fluorescence during the interstimulus period does not differ between WT and Arc KO mice at any age. **B.** Mean ΔF/F amplitudes of visual responses to the maximally exciting visual stimulus for P30 WT and Arc KO mice. Responses are broken down for each age for neurons that respond only to the contralateral eye (Contra), only to the ipsilateral eye (Ipsi), and for binocular neurons broken down further to the mean maximal response to the contralateral eye (Binoc Contra) and ipsilateral eye (Binoc Ipsi). WT and Arc KO response amplitude does not differ at P30 in any of these eye specific responses.

**Supplementary Figure 3 related to figure 5.**
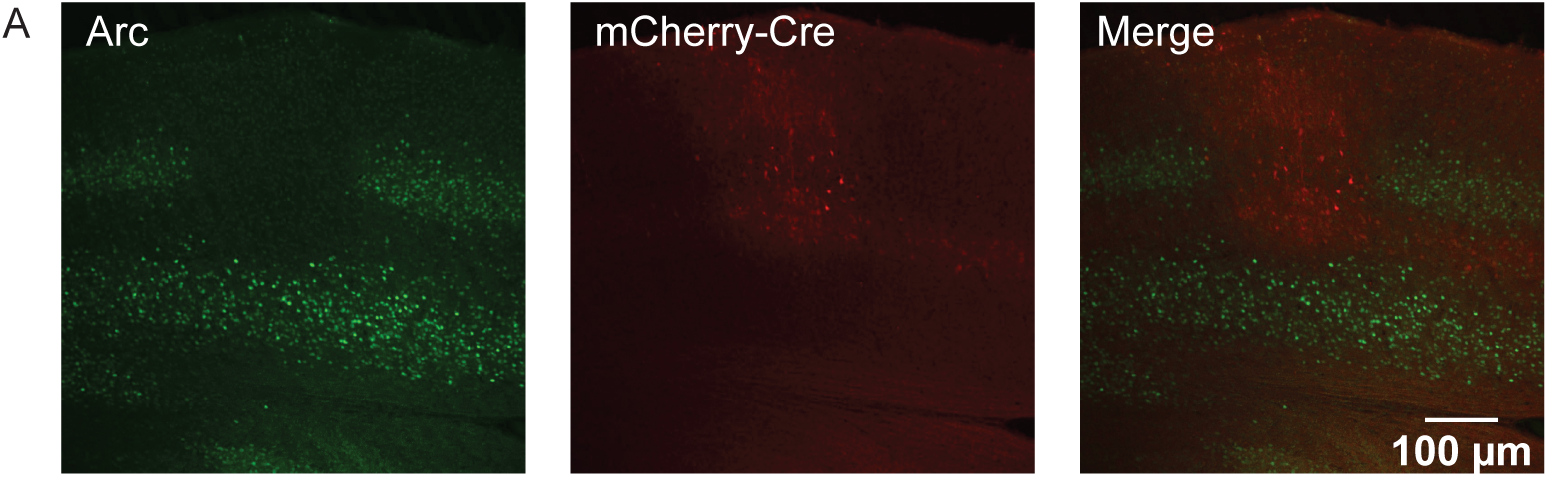
Conditional deletion of Arc expression in binocular visual cortex. **A.** Immunohistochemistry performed in a Cre injected Arc cKO mouse. Arc protein staining (green, left panel) is selectively decreased in the region of mCherry expression (red, middle panel. Right panel, merge).

**Supplementary Table 1:** Response properties of all groups used in the study, separated for monocular (upper) and binocular (lower) cells. Number in parentheses next to each group reports N, number of mice in group. Number listed for response properties reports mean of group ± the standard error of the mean.

| Monocular |  |  |  |  |  |  |
| --- | --- | --- | --- | --- | --- | --- |
| Genotype/Age (N) | Contra OSI | Ipsi OSI | Contra DSI | Ipsi DSI | Contra Pref (°) | Ipsi Pref (°) |
| WT P14 (5) | 0.45±0.05 | 0.61±0.09 | 0.50±0.03 | 0.46±0.07 | 50.9±5.7 | 54.5±5.2 |
| WT P20 (5) | 0.75±0.02 | 0.60±0.08 | 0.37±0.03 | 0.35±0.03 | 40.8±3.5 | 44.6±3.9 |
| WT P30 (6) | 0.73±0.02 | 0.69±0.02 | 0.39±0.02 | 0.36±0.03 | 39.9±2.2 | 42.2±2.2 |
| Arc KO P14 (6) | 0.55±0.05 | 0.48±0.08 | 0.49±0.02 | 0.41±0.03 | 50.9±3.5 | 44.9±4.0 |
| Arc KO P20 (3) | 0.73±0.05 | 0.56±0.06 | 0.40±0.02 | 0.40±0.06 | 40.3±2.1 | 39.1±9.6 |
| Arc KO P30 (4) | 0.73±0.05 | 0.67±0.02 | 0.32±0.05 | 0.34±0.04 | 41.7±2.1 | 48.4±3.7 |
| NR WT (7) | 0.75±0.02 | 0.58±0.04 | 0.33±0.02 | 0.35±0.03 | 42.2±1.2 | 32.2±4.9 |
| DR WT (4) | 0.68±0.05 | 0.67±0.04 | 0.60±0.05 | 0.43±0.04 | 45.6±0.73 | 47.4±5.1 |
| Control cKO P180 (5) | 0.78±0.01 | 0.68±0.06 | 0.37±0.06 | 0.34±0.05 | 39.3±3.5 | 37.4±10.5 |
| Cre cKO P180 (5) | 0.78±0.01 | 0.68±0.03 | 0.34±0.03 | 0.32±0.02 | 38.2±1.7 | 45.0±1.2 |

**Supplementary Table 1:** Response properties of all groups used in the study, separated for monocular (upper) and binocular (lower) cells.
| Binocular |  |  |  |  |  |  |  |
| --- | --- | --- | --- | --- | --- | --- | --- |
| Genotype/Age (N) | Contra OSI | Ipsi OSI | Contra DSI | Ipsi DSI | Contra Pref (°) | Ipsi Pref (°) | ODI |
| WT P14 (4) | 0.44±0.08 | 0.68±0.12 | 0.45±0.04 | 0.45±0.12 | 69.0±6.3 | 78.6±7.2 | 0.39±0.08 |
| WT P20 (5) | 0.72±0.04 | 0.75±0.05 | 0.28±0.04 | 0.32±0.04 | 33.0±2.3 | 32.6±4.5 | 0.45±0.04 |
| WT P30 (6) | 0.71±0.04 | 0.63±0.04 | 0.28±0.03 | 0.30±0.02 | 29.±2.7 | 34.2±4.3 | 0.38±0.03 |
| Arc KO P14 (3) | 0.46±0.23 | 0.51±0.22 | 0.46±0.03 | 0.56±0.10 | 49.5±12.1 | 36.5±12.6 | 0.33±0.07 |
| Arc KO P20 (3) | 0.80±0.04 | 0.76±0.07 | 0.37±0.05 | 0.36±0.14 | 19.6±2.5 | 17.3±9.1 | 0.38±0.03 |
| Arc KO P30 (4) | 0.71±0.05 | 0.70±0.05 | 0.23±0.01 | 0.27±0.01 | 29.8±4.8 | 30.1±5.3 | 0.41±0.02 |
| NR WT (7) | 0.71±0.05 | 0.63±0.08 | 0.23±0.02 | 0.36±0.03 | 34.8±3.7 | 35.3±3.3 | 0.31±0.02 |
| DR WT (4) | 0.59±0.08 | 0.55±0.06 | 0.39±0.09 | 0.27±0.02 | 51.5±8.8 | 44.5±1.5 | 0.31±0.03 |
| Control cKO P180 (5) | 0.81±0.02 | 0.73±0.01 | 0.24±0.05 | 0.26±0.03 | 29.7±5.7 | 27.6±4.7 | 0.41±0.02 |
| Cre cKO P180 (5) | 0.74±0.04 | 0.66±0.06 | 0.26±0.02 | 0.22±0.02 | 36.9±2.3 | 41.1±2.0 | 0.35±0.03 |

